# Decoding the single-cell landscape and intercellular crosstalk in the transplanted liver: a 4-dimension mouse model

**DOI:** 10.1101/2021.01.06.425562

**Authors:** Haitao Huang, Xueyou Zhang, Hui Chen, Shi Feng, Cheng Zhang, Ruihan Chen, Yimou Lin, Qinghua Ji, Qi Ling

**Author notes:** Correspondence and request for materials should be addressed to Qi Ling.

## Abstract

Graft remodeling after transplantation maintains graft functionality and determines graft survival. However, a comprehensive understanding of cellular diversity and interplay during graft remodeling remains to be fully characterized. In this study, we established a well tolerant C57BL/6 to C57BL/6 orthotopic liver transplantation (LT) mice model and observed two stages of graft recovery including an acute phase and a steady phase. We next performed single-cell RNA sequencing (scRNA-seq) and cytometry by time-of-flight (CyTOF) and recorded the cellular hierarchy in the transplanted liver during the two stages. Besides the dynamic change of cell proportion, it was notable that recipient-derived cells took over the transplanted liver in most cell types (e.g., B cells, T cells, dendritic cells, granulocytes and monocytes) except CD206^+^ MerTK^+^ macrophages and CD161^+^ CD49a^+^ CD49b^−^natural killer cells. We then focused on macrophages and captured 5 distinct transcriptional signatures to define novel subclusters. Using a ligand-receptor interaction strategy, we identified specific macrophage-hepatocyte interactions during the acute and stable phases, causing metabolic remodeling in the transplanted liver. Our results delineated a 4-dimension cell atlas (type-proportion-source-time) of the transplanted liver, which sheds light on the physiological process of liver graft maintenance and graft-recipient crosstalk.

## INTRODUCTION

Liver transplantation (LT) is the most effective salvage treatment for various end-stage liver diseases [1]. With the development of the transplant technique and immunosuppressant in the last decades, LT has become a routine surgical procedure with a high success rate [2]. Long-term survival and improved quality of life are the current purposes after LT [3, 4]. In liver recipients, long-term graft survival mostly relies on the prompt recovery of graft function and avoidance of post-transplant complications such as graft dysfunction, rejection, biliary complications and metabolic disorders [5–7].

Graft remodeling has been considered a complex and well-orchestrated process that involves the cellular composition and functional state from multiple lineages [8]. In LT, immune cells derived from the recipient’s peripheral blood invade through the hepatic sinusoid and move towards the interstitial space, where interplay with graft-derived cells to further reorganize immune and metabolic homeostasis [9, 10]. During the early post-LT period, parameters reflecting liver injury and inflammatory response would be predictors for prognosis. For example, alanine aminotransferase (ALT) and γ-GT levels on postoperative day 1 have been considered as independent predictors of early allograft dysfunction [11]. A combined detection of IL-10, IL-17, and CXCL10 hold an excellent prediction property for acute rejection [12]. Therefore, a deep understanding of graft remodeling including the recipient-graft interplay is essential to promote the recovery of the transplanted liver and prevent post-transplant complications.

The single-cell approach has emerged as a powerful tool to deconstruct the composition and function of complex tissues at single-cell resolution [13]. Recently, single-cell RNA sequencing (scRNA-seq) has been applied to dissect the complex landscapes of organogenesis, including liver development [14] and decidualization [15]. The advent of the high-throughput single-cell profiling technology provides a foundation for “transplanted liver atlas”. In this study, we employed scRNA-seq and cytometry by time-of-flight (CyTOF) to comprehensively reveal the cell states and sources involved in liver graft remodeling. We then uncovered an interplay between hepatocytes and macrophages using ligand-receptor interactions based on the transcriptome data [16]. Our analyses constructed a detailed molecular and cellular map and revealed complex intercellular crosstalk in the transplanted liver.

## MATERIALS AND METHODS

### Animals treatments

Male 8-12 weeks old wild-type C57BL/6 mice and C57BL/6-Tg (CAG-EGFP) mice were purchased from Shanghai Model Organisms Center. All mice were housed at 20□ with a 12:12 light-dark cycle in a specific pathogen-free animal facility and fed ad libitum with a normal chow diet. All animal experimental protocols were approved by the Institutional Animal Care and Use Committee and the Ethics Committee of Zhejiang University. Livers from C57BL/6-Tg (CAG-EGFP) mice (all tissue expressing eGFP) used as donors and wild-type C57BL/6 mice were used as recipients for the orthotopic LT model. Surgery was performed as we described before [17]. After the postoperative restoration of temperature and rehydration, mice were sent to the individually ventilated cages for housing. The transplanted liver and serum samples were collected at 3H, 6H,12H, 1D, 3D, 7D, and 14D post-LT. Biochemical parameters including ALT and aspartate aminotransferase (AST) were measured with a BioVision system. The severity of liver ischemia-reperfusion injury (IRI) was graded by modified Suzuki’s histological criteria [18].

### Immunohistochemistry

Immunohistochemistry was performed as we described previously [19]. Liver specimens (4μm) were stained with hematoxylin and eosin (H&E). Two independent pathologists reviewed and confirmed the pathohistological features of all the samples. For staining of neutrophils and macrophages, anti-Ly6g (Abcam, USA, ab238132) and anti-F4/80 (Bio-rad, USA, MCA497R) were used. For terminal deoxynucleotidyl transferase-mediated dUTP nick-end labeling (TUNEL) staining, liver specimens were stained using the In Situ Cell Death Detection Kit (Roche, USA, 11684817910) according to the manufacturer’s protocol. Labeled cells were counted with 5 randomly selected 400× fields from each sample.

### Quantitative real-time PCR

Total RNA was extracted using TRIzol reagent (Invitrogen, USA, 15596018) and quantitative real-time PCR was performed using the HiScript II One Step qRT-PCR SYBR Green Kit (Vazyme Biotech, China, Q221-01) in an Applied Biosystems QuantStudio 5 Real-Time PCR Instrument (Applied Biosystem, USA) as we described previously [17]. The detailed primer information was provided in Table S1.

### Tissue dissociation and preparation

Liver tissues were obtained from pre- and post-LT period. The tissues were transferred into sterile RNase-free culture dishes containing an appropriate amount of calcium-free and magnesium-free PBS on ice and cut into 0.5 mm^3^ pieces. Then the tissues were washed with PBS and dissociated into single cells in dissociation solution (0.35% collagenase IV, 2 mg/ml papain, 120 Units/ml DNase I) in 37□ water baths with shaking at 100 rpm for 20 min. Digestion was terminated by PBS containing 10% fetal bovine serum (V/V). The suspensions were passed through 100 μm cell sieves (Corning, Japan, 431752) and then 40 μm cell sieves (Corning, Japan, 431750). The flow-through was centrifuged and resuspended in 5 ml of red blood cell lysis buffer (Invitrogen, USA, 00-4300) for 10 min. After incubation, the suspension was centrifuged at 300 g for 5 min at 4□. For scRNA-seq, the overall cell viability was confirmed by trypan blue (over 85%), single-cell suspensions were counted using a Countess II Automated Cell Counter (Invitrogen, USA, 00-4300), and concentration adjusted to 700-1200 cells/μl.

### CyTOF

Liver single-cell suspensions were processed for CyTOF analysis according to the standard protocol as we described previously [17]. Briefly, the single-cell suspensions were labeled by 194Pt for 5 min to distinguish live or dead and incubated with Fc blocking mix for 20 min at room temperature to block Fc receptors. Then the single-cell suspensions were stained for 30 min with an antibody mix panel (Table S2), followed by DNA Intercalator-Ir overnight staining. Intracellular markers were stained using an intracellular antibody mix for 30 min. Finally, the single cells were washed with PBS, and the signals were detected using a CyTOF system (Fluidigm, USA). The types of immune cells were identified via t-distributed stochastic neighbor embedding (tSNE), followed by density clustering [20]

### scRNA-seq

Liver single-cell suspensions were loaded into the 10x Chromium Controller, and the 10x Genomics Chromium Single Cell 3’ kit (V3) was applied according to the manufacturer’s protocol. Libraries were sequenced on an Illumina NovaSeq 6000 sequencing system. We used FastQC to perform basic statistics on the quality of the raw reads. Raw reads were demultiplexed and mapped to the reference genome by 10x Genomics Cell Ranger pipeline (https://support.10xgenomics.com/single-cell-geneexpression/software/pipelines/latest/what-is-cell-ranger). All downstream single-cell analyses were performed using Cell Ranger (version 3.1.0) and Seurat (version 3.2.0) unless mentioned specifically. In brief, for each gene and each cell barcode (filtered by CellRanger), unique molecule identifiers were counted to construct digital expression matrices. Secondary filtration by Seurat: A gene with expression in more than 3 cells was considered as expressed, and each cell was required to have at least 200 expressed genes. We removed low-quality cells with > 20% mitochondrial RNA content and included cells with > 200 and < 6000 genes expressed per cell. Seurat was used to normalizing data, dimensionality reduction, clustering and differential expression analysis. To eliminate batch effects, a reference Seurat object was generated by identifying anchor genes between samples as described previously [14]. For clustering, highly variable genes were selected and the principal components based on those genes were used to build a graph. The tSNE analysis was used for dimension reduction and visualization of gene expression. For differential expression analysis, marker genes for each cluster were determined by comparing the cells of each cluster to all other cells using the FindAllMarkers function in Seurat. For all comparisons between groups and clusters, only genes expressed by at least 10% of cells were included. Gene Ontology (GO) enrichment analysis of marker genes was implemented by the clusterProfiler R package (version 3.16.1), in which gene length bias was corrected. GO terms with corrected *P* value less than 0.05 were considered significantly enriched by marker genes. gene set variation analysis (GSVA) analysis of distinct clusters was performed by the GSVA R package (version 1.36.2)[21].

### Cell-cell interplay analysis

To enable a systematic analysis of cell-cell communication, we applied nichenetr package (version 1.0.0) to predict cell-cell interplay as previously reported [22]. NicheNet could predict ligands of sender cells that are the most active in affecting gene expression of receiver cells, through the correlation of ligand activity with genes characterized as targets in a prior model that integrates existing knowledge on ligand-to-target signaling path [16]. Using macrophages and hepatocytes subsets, we applied NicheNet to predict ligands that might influence transcription during hepatocytes metabolic remodeling. In the integration analysis, hepatocytes were defined as the receiver population and all macrophage subclusters were defined as potential sender cells. In separate analyses, target macrophage subclusters were defined as potential sender cells. Gene sets of interest were defined based on metabolic-related genes in MSigDB gene sets, including regulation of glucose metabolic process, insulin signaling pathway and lipid homeostasis (Table.S3). A background gene set includes all other genes expressed in at least 10% of cells in hepatocytes. Activities of ligands were ranked by calculating Pearson correlation coefficients of the ligand-target regulatory potential scores for each ligand and the target indicator vector, which defines a gene as present or absent within the gene set of interest. The circlize R package (version 0.4.10) was used to represent the regulatory networks between the predicted ligands in distinct macrophage subclusters and their receptors and targets expressed in hepatocytes. Cytoscape (version 3.8.0) was used to depict the regulatory networks between the predicted ligands in the target macrophage subclusters and their targets expressed in hepatocytes.

## RESULTS

### Liver graft recovery following transplantation

Liver graft injury is inevitable due to multiple factors including IRI and surgical stress. To monitor the liver graft recovery after transplantation, we established an orthotopic LT mice model and collected serum and liver tissue samples during the perioperative period (Pre-LT, 3H, 6H, 12H, 1D, 3D, 7D and 14D post-LT; n = 3/group, Fig.1A). We observed sharply increased ALT and AST levels at 3H post-LT. From 3H to 1D, serum ALT and AST levels gradually decreased and maintained steady after 1D (Fig.1B). We next assessed the morphology organization of transplanted livers using histopathology methods. H&E staining showed the destruction of hepatic architecture as presented by sinusoidal congestion, hepatocyte necrosis, and ballooning degeneration. Suzuki’s histological grading of hepatocellular damage displayed the highest score at 1D (Fig.1C). We further found consistent results in hepatocellular necrosis/apoptosis assessment (Fig.1D). Moreover, we assessed inflammatory cell infiltration using immunohistology and observed that neutrophils and macrophages maintained at high levels during the first day and reduced thereafter (Fig.1E). The cytokine and endothelial cell activation marker expression patterns showed a similar trend with inflammatory cells (Fig.1F). The results suggested an acute injury of graft function within 1D (acute phase) and a steady recovery of graft function after 3D (stable phase).

**Figure 1.**
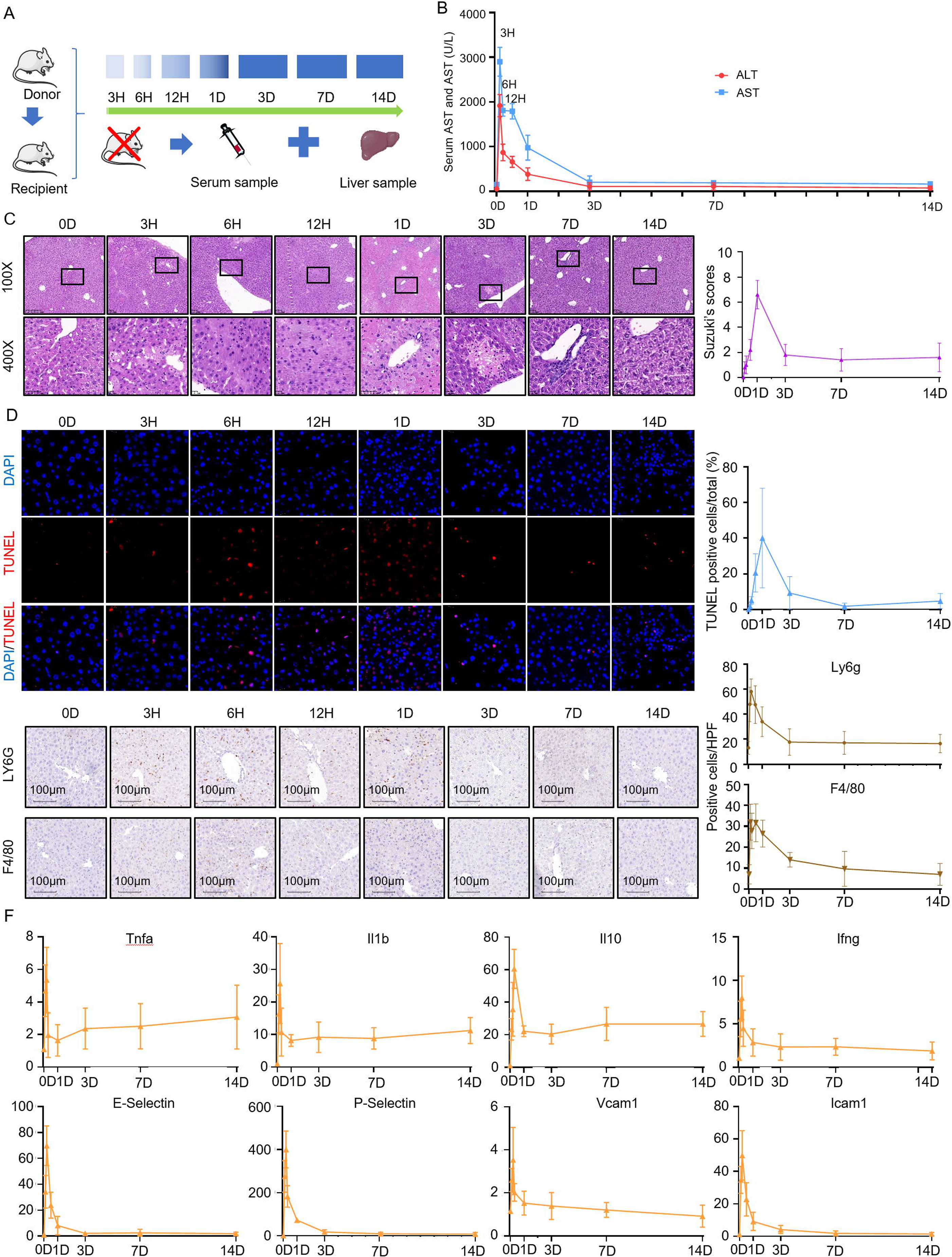
Liver graft recovery in orthotopic liver transplantation mice model. (A) Experimental design in mice. (B) Serum ALT and AST levels in mice. (C) H&E staining and Suzuki’s histological grading of transplanted livers. (D) TUNEL staining of transplanted livers. (E) Neutrophils (LY6G+) and macrophages (F4/80+) infiltration in transplanted livers. (F) The expression of cytokine and endothelial cell activation markers in transplanted livers. ALT, alanine aminotransferase; AST, aspartate aminotransferase; H&E, hematoxylin and eosin; TUNEL, terminal deoxynucleotidyl transferase-mediated dUTP nick-end labeling.

### Mapping cellular hierarchy in the transplanted liver

To map the cellular diversity and the dynamic change during the perioperative period, we generated scRNA-seq profiles from 6 liver samples (pre-LT, 3H, 6H, 12H, 3D and 7D post-LT) using 10x Genomics sequencing (Fig.2A). We obtained a total of 22,119 single-cell transcriptomes for analysis. We conducted gene expression normalization, principal component analysis (PCA), and tSNE. Finally, all liver cells could be assigned to 10 distinct types based on marker gene expression and scMCA (http://bis.zju.edu.cn/MCA/blast.html). They were granulocyte (38.8%, marked with *Ly6g*), T cells (14.5%, marked with *Cd3d*), macrophages (13.1%, marked with *Adgre1*), endothelial cells (9.4%, marked with *Bmp2*), B cells (8.3%, marked with *Cd79a*), dendritic cells (DCs, 6.4%, marked with *Itgax*), natural killer (NK) cells (5.9%, marked with *Klrb1c*), hepatocytes (2.1%, marked with *Apoa1*), biliary epithelial cells (1.0%, marked with *Krt19*) and fibroblast (0.4%, marked with *Acta2*, Fig. 2B). The differentially expressed genes (DEGs) and GO enrichment analysis further confirmed the cell identity (Fig. 2C and Fig.S1A). Moreover, we observed a significant variation in the proportion of immune cells during the perioperative period (Fig.2D). For instance, granulocytes increased rapidly from 8.1% to 78.3% at 6H post-LT and macrophages increased from 9.7% to 29.7% at 12H post-LT, indicating an IRI-induced inflammatory response. Both T cells and B cells dropped to less than 15% after 6H post-LT, suggesting a tolerance in our model. (Fig.2D and Fig.S1B).

**Figure 2.**
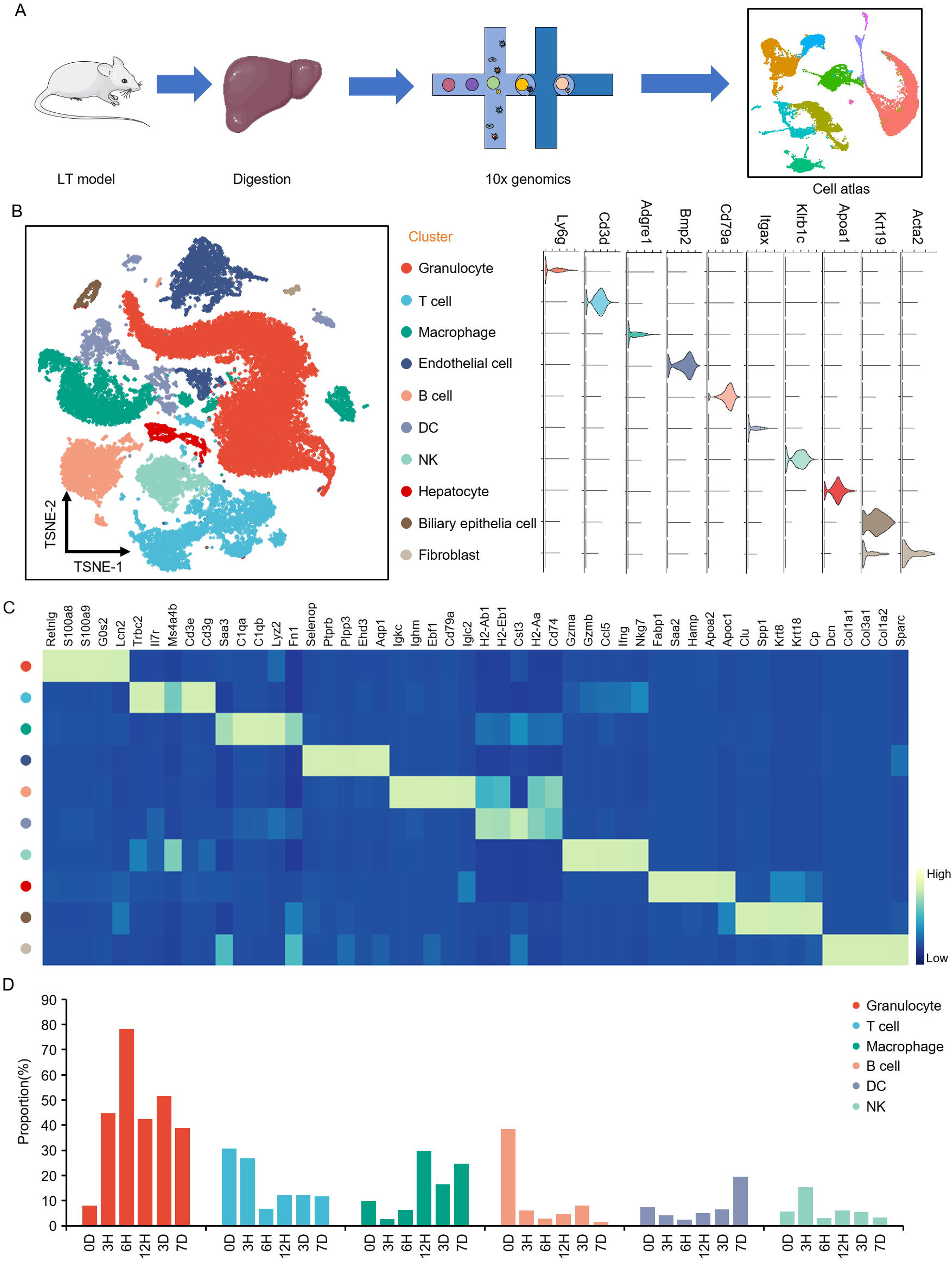
The cellular hierarchy in the transplanted liver. (A) Schematic diagram of the scRNA-seq analysis workflow. (B) tSNE plots for the cell type identification of single cells (left); Violin plots showing marker genes for 10 distinct cell types (right). (C) Heatmap showing the top 5 DEGs in each cell type. (D) Bar plots showing the proportion of immune cells at pre-LT, 3H, 6H, 12H, 3D, and7D post-LT. scRNA-seq, single-cell RNA sequencing; tSNE, t-distributed stochastic neighbor embedding; DEGs, differentially expressed genes; LT, liver transplantation

### The cellular source in the transplanted liver

To explore the comprehensive graft-recipient interaction following LT, we used an eGFP-labeled donor mouse to distinguish graft-derived cells from recipient-derived cells. The efficiency of eGFP fluorescence was 90% according our previous report [17]. We acquired liver tissue samples at pre-LT, 12H, 1D, 3D, 7D and 14D post-LT and purified CD45^+^ liver nonparenchymal cells for CyTOF analysis (Fig. 3A), which comprises 42 antibodies. Cells were initially gated and t-SNE visualized to distinct populations. A schematic clustering explanation is depicted in Fig. 3B. Among the 29 clusters, No. 5 could not be categorized and the other 28 were defined as 7 cell types including B cells (2 clusters, marked with CD19), T cells (5 clusters, marked with CD3), DCs (5 clusters, marked with CD11c), NK cells (3 clusters, marked with NKp46 or CD49b), macrophages (8 clusters, marked with F4/80), granulocytes (2 clusters, marked with LY6G) and monocytes (3 clusters, marked with LY6C; Fig. 3C). We observed that graft-derived cells (GDCs) dramatically decreased from 100% to 11%, whereas the recipient-derived cells (RDCs) sharply increased from 0% to 89% within the first day post-LT (Fig. 3D). The cell source maintained a steady level from 1D to 14D. RDCs accounted for most cells at 14D post-LT (26/29 clusters) (Fig. 3E). Of note, two M2 macrophage clusters (cluster 1 and cluster 23; F4/80^+^ CD206^+^ MerTK^+^) and one NK cell cluster (cluster 16; CD161^+^ CD49a^+^ CD49b^−^) were composed of around half GDCs at the stable phase (Fig. 3F).

**Figure 3.**
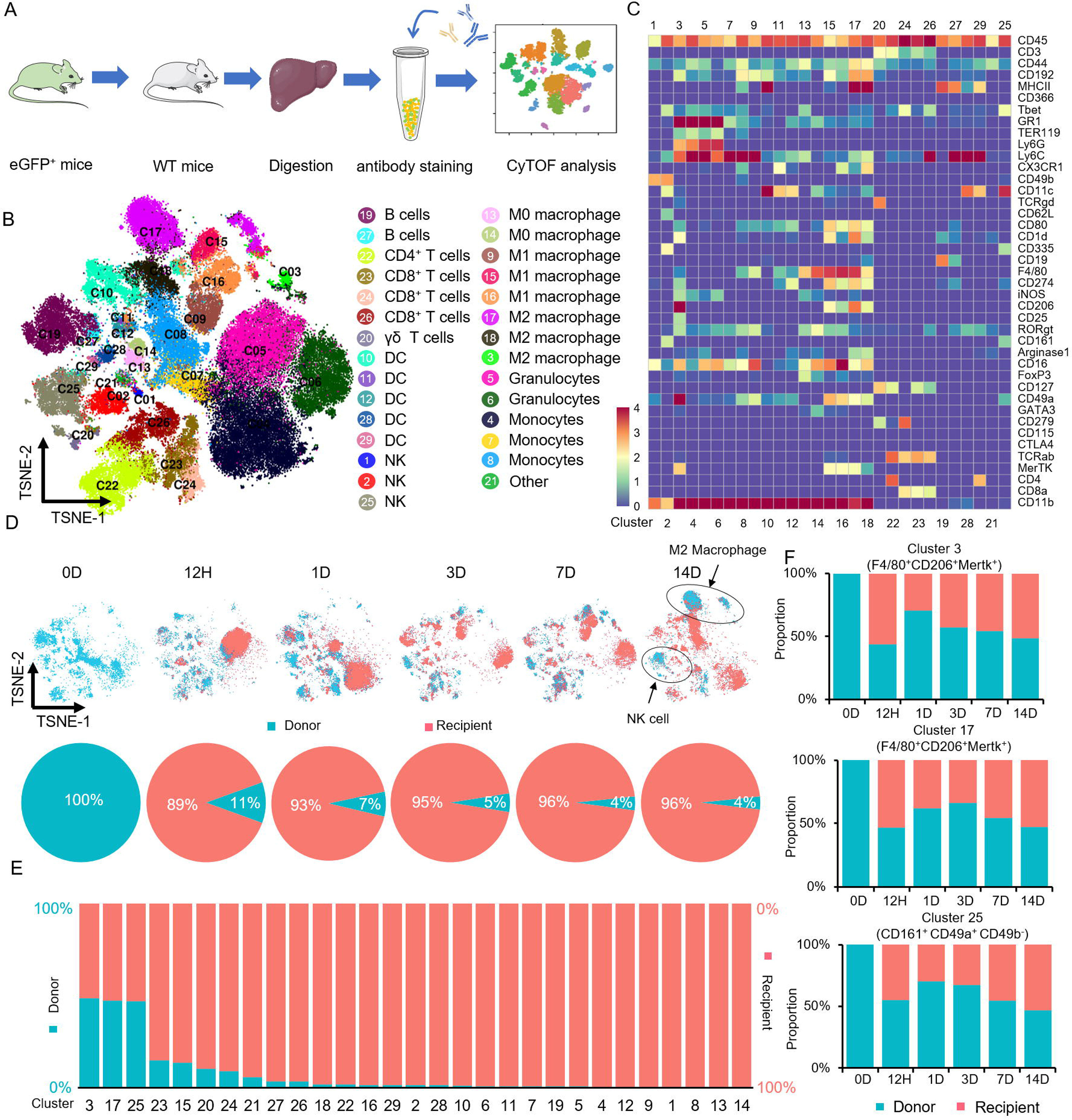
The cellular source in the transplanted liver. (A) Schematic diagram of the CyTOF analysis workflow. (B) tSNE plots for the cell type identification of single cells. (C) Heatmap showing the expression levels of markers in distinct clusters. (D) tSNE and pie plots showing the proportion of GDCs and RDCs at pre-LT, 12H, 1D, 3D, 7D and 14D post-LT. (E) Bar plots showing the proportion of GDCs and RDCs in distinct clusters at 14D post-LT. (F) Bar plots showing the proportion of GDCs and RDCs in cluster 1, 23 and 16 at pre-LT, 12H, 1D, 3D, 7D and 14D post-LT. CyTOF, cytometry by time-of-flight; tSNE, t-distributed stochastic neighbor embedding; GDCs, graft-derived cells; RDCs, recipient-derived cells; LT, liver transplantation

### Metabolic remodeling in the transplanted liver

Considering the specific source and vital role of the macrophage in hepatic homeostasis particularly glucose and lipid metabolism, we focused on the metabolic remodeling by analyzing the macrophage-hepatocyte interplay. First, we re-analyzed the scRNA-seq data of macrophages and stratified them into 5 subclusters with distinct transcriptomic signatures (Fig. 4A). We identified the function of different subclusters using GSVA, which showed that subclusters 0, 1, 2, 3 and 4 were associated with pro-inflammation, ribosome and proteasome, metabolism, immune regulation and allograft rejection, respectively (Fig. 4B). We found that subcluster 0 and 1 expressed high levels of *Ly6c2* and *Ccr2* (immature monocyte-derived macrophages genes), while subcluster 3 expressed low level *Ccr2* and high level of *Cx3cr1* (mature monocyte-derived macrophages genes). Subcluster 2 expressed a certain number of *Adgre1*, *Clec4f* and *Timd4* (Kupffer cells genes). Moreover, we assumed that subcluster 0 and 1 belonged to the M1 phenotype due to the pro-inflammatory markers such as *Tnf*, *Ccl2* and *Ccl9*, while subcluster 2 belonged to the M2 phenotype due to the prominent expression of M2 markers such as *Mrc1* (coding CD206), *Cd163* and *Mertk* (Fig. 4C). We observed the dynamic change of different subclusters in the acute and stable phases. For instance, subcluster 0 sharply increased in the acute phase and decreased in the stable phase, whereas subcluster 2 dramatically decreased in the acute phase and partially recovered in the stable phase (Fig. 4D).

**Figure 4.**
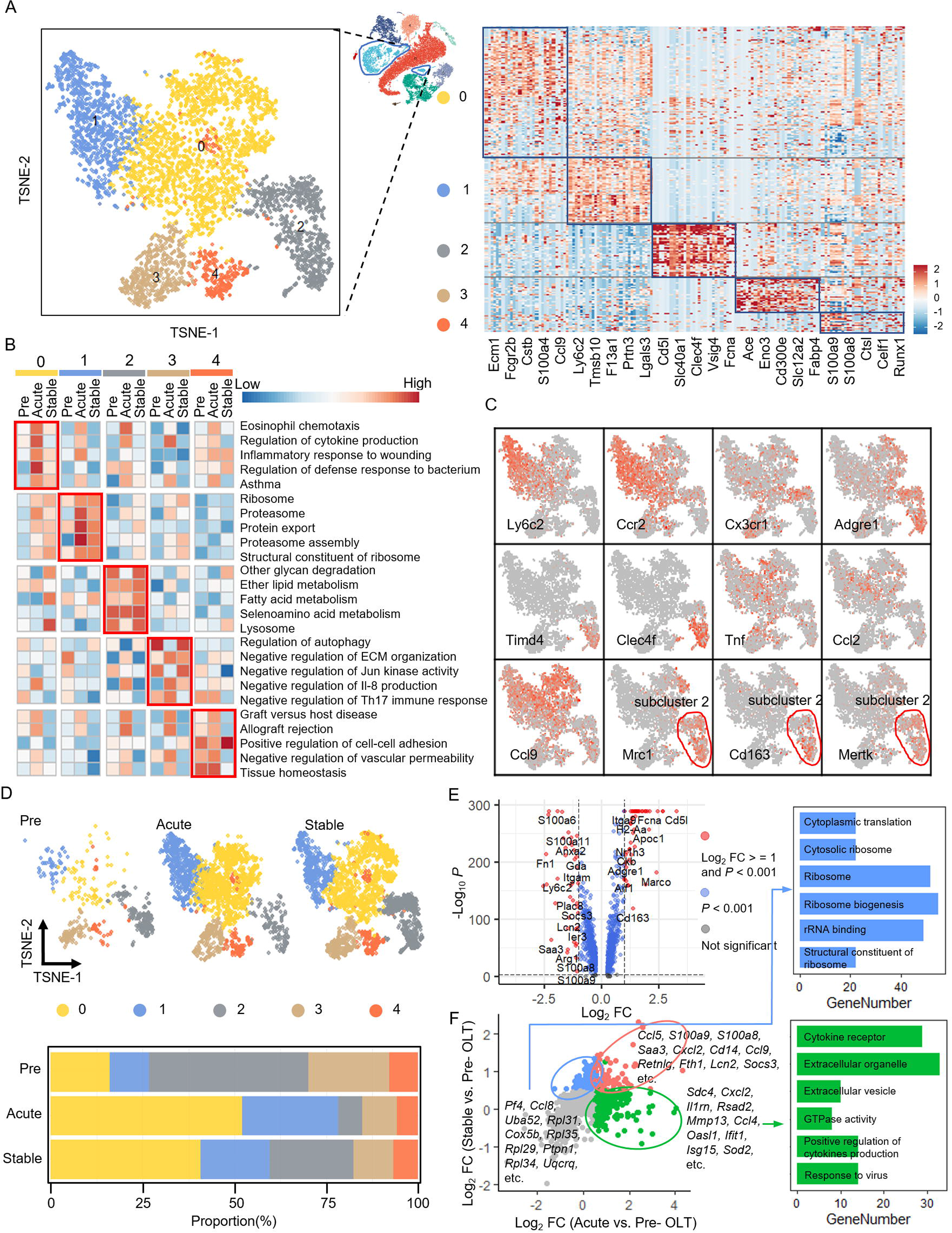
Distinct macrophage subclusters in the transplanted liver. (A) tSNE plots of total macrophages (left); Heatmap showing the top 30 DEGs for each subcluster (right). (B) Pathway activity (scored by GSVA) in 5 subclusters. (C) tSNE plots color-coded for the expression (gray to red) of marker genes for the distinct subclusters. The red circle showed subcluster 2. (D) tSNE and bar plots showing the proportion of distinct subclusters at pre-LT, acute phase and stable phase. (E) Volcano plot exhibiting DEGs between subcluster 2 and other macrophages. (F) Scatterplots showing genes upregulated in either acute phase and/or stable phase of subcluster 2 (left); Bar plots showing enriched GO terms of the genes overexpressed by subcluster 2 in the acute phase and stable phase (right). tSNE, t-distributed stochastic neighbor embedding; DEGs, differentially expressed genes; GSVA, gene set variation analysis; LT, liver transplantation; GO, Gene Ontology.

We should also note that subcluster 2 was comprised of both graft- and recipient-derived cells (F4/80^+^ CD206^+^ MerTK^+^ Macrophages) according to the CyTOF findings. In subcluster 2, the genes involved in anti-fibrosis and metabolism such as *Cd5l* [23], *Timd4* [24], *Clec2* [25] and *Pltp* [26] were significantly up-regulated and the genes related to inflammation such as *Tnf*, *Il1b* and *Cxcl2* [27] were significantly down-regulated compared to other macrophages (Fig. 4E). Furthermore, a comparison of subcluster 2 in the acute and stable phases with that at pre-LT revealed distinct transcriptional programs (Fig. 4F). Compared with pre-LT, genes highly expressed in subcluster 2 in the acute phase were enriched in cytokine receptors and extracellular vesicles. In the stable phase, highly expressed genes were mainly enriched in cytoplasmic translation and ribosome.

Second, we analyzed the metabolic activity of hepatocytes using GSVA and found decreased gluconeogenesis, insulin signaling and lipid metabolic process whereas increased glycolysis in the acute phase. The changed glucose and lipid metabolism well recovered in the stable phase (Fig.5A). Next, we performed NicheNet, which could show ligand-receptor interactions and ligand-to-target signaling paths [16], to analyze the macrophage-hepatocyte interplay. A circle plot was designed to show the interaction between the receptors and target genes in hepatocytes and the most active ligands in macrophages (Fig.5B). Macrophage subcluster 0, 1, 2 and 4 presented potential specific ligands that regulate insulin signal, glucose and lipid metabolic genes in hepatocytes. For example, *Ceacam1*, *Cxcl2*, *Tnf* and *Itgam* were most expressed in subcluster 0, while *Cxcl13*, *Cxcl16*, *Apoe* and *Nampt* were expressed in subcluster 2.

**Figure 5.**
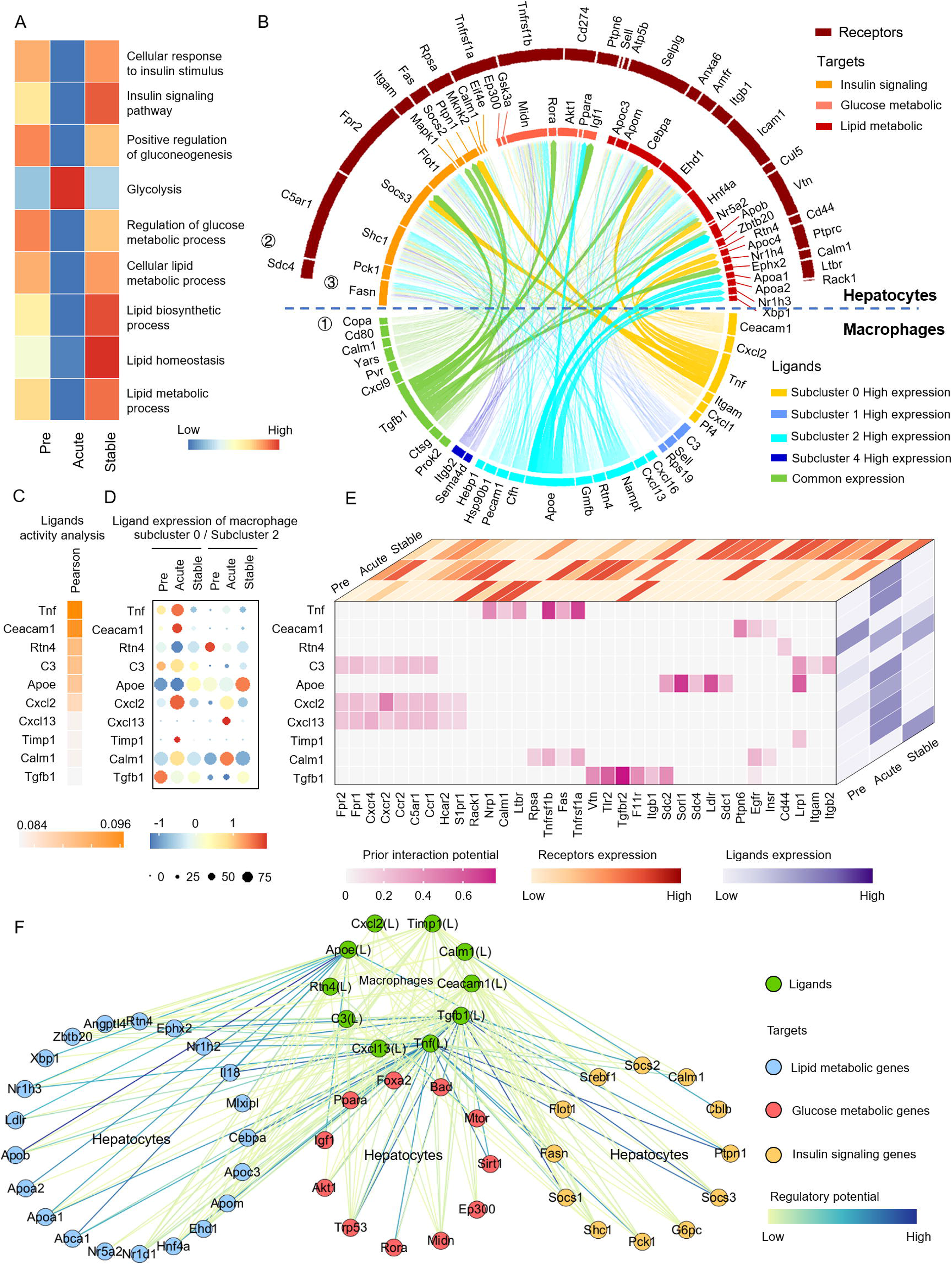
Macrophage-hepatocyte interplay in the transplanted liver. (A) Heatmap showing the metabolic-related pathway activity (scored by GSVA) in hepatocytes. (B) Circle plot showing links between ① predicted ligands from subcluster 0, subcluster 1, subcluster 2, subcluster 4 or common, ② their associated receptors found on hepatocytes and ③ metabolic genes potentially targeted by the ligand-receptor pairs. (C) Potential upstream ligands in subcluster 0 and subcluster 2; (D) Bubble plot showing the expression of potential upstream ligands in subcluster 0 and subcluster 2 at pre-LT, acute phase and stable phase; (E) Cube plot showing the potential receptors expressed by hepatocytes associated with each potential ligands, and their expression at pre-LT, acute phase and stable phase. (F) Network plot showing links between predicted ligands and metabolic genes. GSVA, gene set variation analysis; LT, liver transplantation.

Based on the abundant crosstalk between subcluster 0 / 2 and hepatocytes, we isolated the 3 populations and further predicted the specific ligand-receptor interactions. Ligands activity analysis showed the top 10 ligands in subcluster 0 and 2 (Fig.5C). For instance, *Tnf*, coding cytokine involved in hepatocyte cell death as well as insulin resistance and lipid accumulation [28, 29], was mainly expressed by subcluster 0. The expression of *Tnf* increased significantly in the acute phase and recovered in the stable phase (Fig.5D). In addition, hepatocytes strongly expressed Tnf receptor genes *tnfrsf1a* and *tnfrsf1b* in the acute phase, suggesting an enhanced Tnf-Tnf receptor interaction in the acute phase (Fig.5E). Target genes prediction showed that *Tnf* could regulate different types of metabolic genes, including glucose metabolism genes (*Sirt1*, *Ppara*, *Akt1*, etc.), lipid metabolism genes (*Cebpa*, *Nr1h2*, *Apoa1*, etc.) and insulin pathway genes (*Pck1*, *Socs3*, *Ptprt1*, etc.; Fig.5F). For another example, *Apoe*, coding apolipoprotein E that involved in lipid and lipoprotein metabolism, was mainly expressed by subcluster 2. The expression of *Apoe* and Apoe receptor genes, such as *Sorl1* and *Ldlr*, decreased in the acute phase and increased in the stable phase, suggesting an enhanced Apoe-Apoe receptor interaction in the stable phase. Furthermore, target genes prediction showed that *Apoe* mainly regulated lipid metabolism-related genes in hepatocytes such as *Srebf1*, *Abca1*, *Apoa1*, *Apoa2*, *Apob*, *Igf1*, *Ldlr*, etc. Other ligands such as *Ceacam1*, *Rtn4*, *C3*, *Cxcl2*, *Cxcl13*, *Timp1*, *Calm1* and *Tgfb1*, were also predicted to regulate a series of metabolic genes in hepatocytes and ultimately contribute to the changes in metabolism-related genes during the perioperative period.

## DISCUSSION

The occurrence of primary nonfunction, graft dysfunction or rejection dramatically reduced both graft and recipient survival. Therefore, graft recovery during the 7-10 days after LT greatly determines the outcome [30]. In this study, we established a tolerant mice model to understand the physiological process of graft recovery and graft-recipient interaction during the early post-LT period. According to the dynamically detected biochemical and pathological parameters, we proposed a two-phase theory of graft recovery following LT, which includes an acute phase and a stable phase. Acute phase was represented by sharply increased hepatocyte damage and inflammatory cell infiltration, occurring at very early stage typically within 24h following LT. The emergence of the peaks of aminotransferases in our model was consistent with most of previous studies [18, 31]. Also, the graft injury would be affected by multiple factors such as ischemia time and perfusate, causing diversity in graft recovery [32, 33]. Stable phase showed a well recovered liver function after the acute graft injury, which probably started from 3D after LT. Then we conducted a single cell technology for the in-depth understanding of graft remodeling in both acute and stable phases.

We described the transcriptomic landscape of the transplanted liver and depicted the cellular atlas, including the cell types and proportion. The diversity of liver parenchymal and non-parenchymal populations was consistent with previous studies both in mice [34] and human [21]. Given the distinct cell types, we found that the cellular composition of the transplanted liver has changed dramatically during the perioperative period. The persistently elevated inflammatory cells (granulocytes and macrophages) in the transplanted liver throughout the acute and stable phases revealed an IRI-induced inflammatory response, which has been associated with the development of post-transplant complications such as tumor recurrence, diabetes, hyperlipidemia and fibrosis [35]. In addition, the transplanted liver in the stable phase had decreased proportions of T cell and B cell, which were known to promote rejection [36], indicating tolerance in the stable phase.

We next focused on the graft-recipient source in distinct cell types. Our previous pioneering study using the same strategy and focusing on the monocyte-derived cells has revealed that RDCs took over the majority of cells, including monocytes, DCs, M0 and M1 macrophages. This study further explored the cellular source of various nonparenchymal (e.g., macrophages, NK cells, DCs, endothelial cells, T cells and B cells) in the transplanted liver. The results were consistent with our previous findings that GDCs disappeared in a short time and RDCs made the major contribution to Kupffer cells pool. The result challenged the classical theory that Kupffer cells are locally proliferate and self-renew [37] at least under the circumstance of LT. Furthermore, not only the monocytes-derived cells, most nonparenchymal cells were refreshed by RDCs at steady phase, suggesting a pivotal role of graft-recipient crosstalk during graft remolding. It was very interesting to find that GDCs were still accounted for a half of proportion in certain cell populations including F4/80^+^ CD206^+^ MerTK^+^ Kupffer cells and CD161^+^ CD49a^+^ CD49b^−^ NK cells. F4/80^+^ CD206^+^ MerTK^+^ Kupffer cells have been associated with the resolution of inflammation and the compromise repair after ischemia-reperfusion [38, 39], as well as the improved hepatic insulin sensitivity [40]. However, these macrophages with an anti-inflammatory and reparative phenotype have been revealed to drive hepatocellular carcinoma progression and metastasis [37]. In addition, a recent study on CD49a^+^ liver-resident NK have showed that a role for CD49a^+^ NK cells in the negative regulation of immune responses and the development of hepatocellular carcinoma[41]. Investigation of diseases associated with Kupffer cells and NK cells after LT (e.g., liver fibrosis [42], rejection[43] and hepatocellular carcinoma [44]) should consider both the graft and recipient genetic backgrounds.

Our previous studies in both large human cohorts [45, 46] and mice models [19] demonstrated an imbalanced hepatic metabolic homeostasis after LT. This study not only proved that hepatocytes showed a metabolic reprogram after LT including insulin resistance, glucose and lipid metabolic homeostasis, but also revealed a comprehensive interplay between macrophage and hepatocytes in the transplanted liver. In previous studies, scRNA-seq analysis has been used to elucidate the diversity of macrophages in human and mouse livers [47, 48]. In this study, we defined 5 macrophage subclusters in the transplanted liver and preliminarily explored their functional transformation at different phases, such as subcluster 0 with high pro-inflammatory activity in the acute phase and subcluster 4 with suppressed rejection activity in the stable phase. Further exploring the interplay between liver macrophages and hepatocytes, macrophages subcluster 0 and subcluster 2 hold the most abundant ligand-receptor pairs and excellent regulatory potential, suggesting an effectively regulation in the metabolic remodeling. For instance, the Tnf-Tnf receptor pair in the acute phase was significantly enhanced, which have been found to be crucial for the development of metabolic disorders [49]. In addition, an enhanced Apoe-Apoe receptor pair, promoting the clearance of remnants of triglyceride-rich lipoproteins[50], was found to contribute to the stable phase. Our scRNA-seq data and functional analysis showed that the metabolic remodeling of the transplanted liver was a very complex process, involving a regulatory network of a series of ligands and receptors between macrophages and hepatocytes.

The study had both advantages and limitations. First, to the best of our knowledge, this is the first study to map a 4-dimension cell atlas of the transplanted liver based on a well tolerant LT mice model. The open source website construction is in progress and the datasets would be available for further analyses such as the comparison with the rejection model or marginal graft model. Second, our finding shed a light on the intrahepatic donor-recipient crosstalk, which is the key to understand liver transplant immunology and pathophysiology. For instance, the F4/80^+^ CD206^+^ MerTK^+^ hepatic macrophages had both donor- and recipient-sources. Since this subclass involves in liver fibrosis, nonalcoholic steatohepatitis and tumor recurrence, both donor and recipient genetic background potentially affect the long-term survival of recipients [37, 51]. The predicted ligand-receptor interactions need to be further proved and more cellular interplays need to be explored.

## AUTHOR CONTRIBUTIONS

Q.L. designed the research; H.H., S.F., T.R., and H.C. performed the research; H.H., X.Z and Q.J. analyzed the data; H.H. and R.C. wrote the manuscript; Q.L. reviewed the manuscript.

## Abbreviations

LT: liver transplantation
scRNA-seq: single-cell RNA sequencing
CyTOF: cytometry by time-of-flight
ALT: alanine aminotransferase
AST: aspartate aminotransferase
IRI: liver ischemia-reperfusion injury
H&E: hematoxylin and eosin
TUNEL: terminal deoxynucleotidyl transferase-mediated dUTP nick-end labeling
tSNE: t-distributed stochastic neighbor embedding
GO: Gene Ontology
GSVA: gene set variation analysis
PCA: principal component analysis
NK: natural killer
DCs: dendritic cells
DEGs: differentially expressed genes
GDCs: graft-derived cells
RDCs: recipient-derived cells

## SUPPLEMENTAL

**Figure S1 Function and proportion of each cell type.** (A) GO analysis of DEGs in distinct cell types. (B) tSNE plots of distinct cell types at pre-LT, 3H, 6H, 12H, 3D, and7D post-LT. GO, Gene Ontology; DEGs, differentially expressed genes; tSNE, t-distributed stochastic neighbor embedding; LT, liver transplantation

**Table S1.**
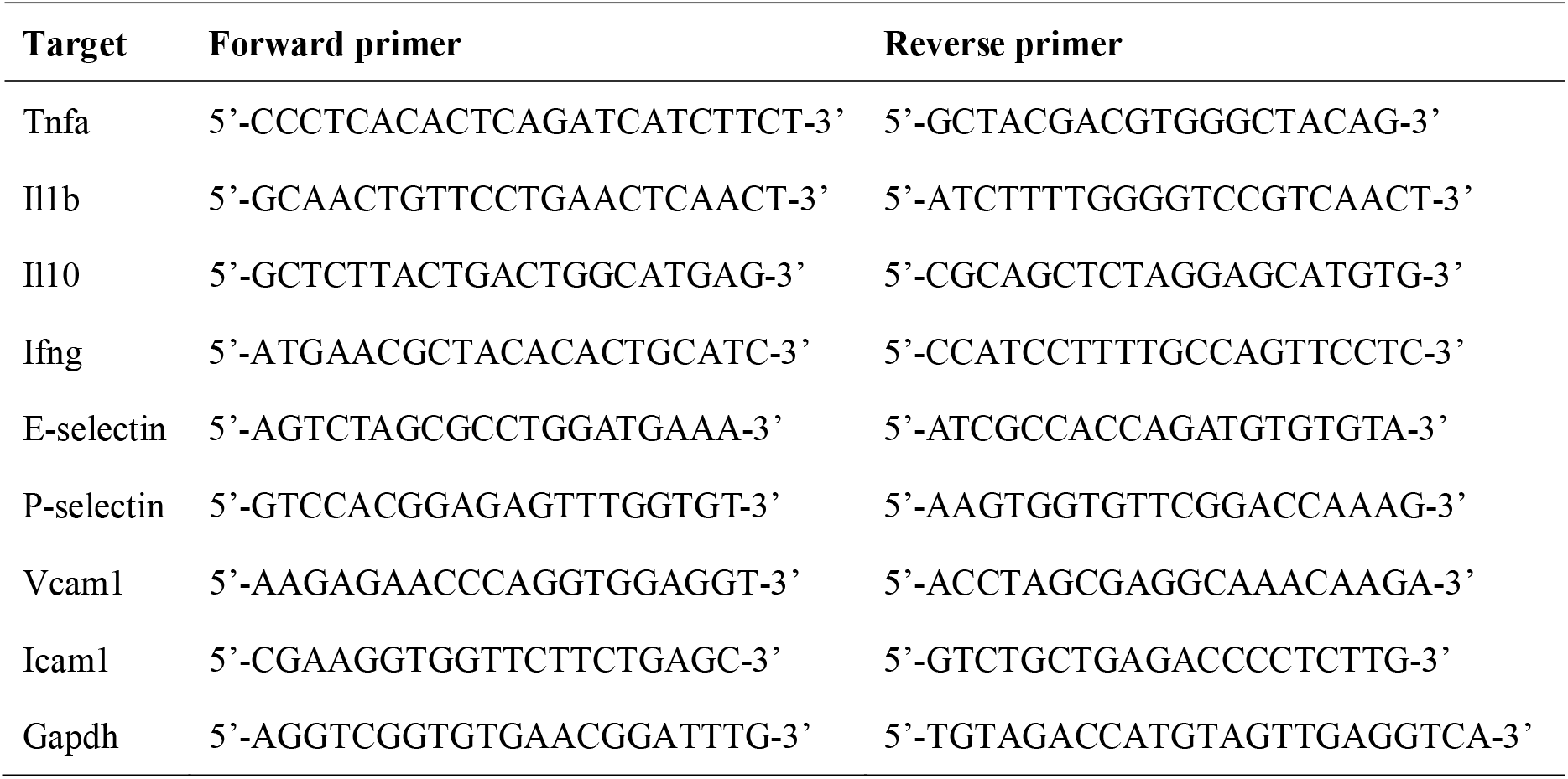
List of primers used.

**Table S2.** Antibody panel.

| Label | Marker | Label | Marker | Label | Marker | Label | Marker | Label | Marker |
| --- | --- | --- | --- | --- | --- | --- | --- | --- | --- |
| 89Y | CD45 | 147Sm | Ly6G | 156Gd | CD1d | 165Ho | CD161 | 174Yb | CTLA4 |
| 115In | CD3 | 148Nd | Ly6C | 157Gd | CD335 | 166Er | Arginase1(Intra) | 175Lu | TCRab |
| 139La | CD44 | 149Sm | CX3CR1 | 158Gd | CD19 | 167Er | CD16 | 176Yb | MerTK |
| 141Pr | CD192 | 150Nd | CD49b | 159Tb | F4/80 | 168Er | FoxP3(Intra) | 197Au | CD4 |
| 142Nd | MHCII | 151Eu | EGFP(Intra) | 160Gd | CD274 | 169Tm | CD127 | 198Pt | CD8a |
| 143Nd | CD366 | 152Sm | CD11c | 161Dy | iNOS(Intra) | 170Er | CD49a | 209Bi | CD11b |
| 144Nd | Tbet(Intra) | 153Eu | TCRgd | 162Dy | CD206(Intra) | 171Yb | GATA3(Intra) |  |  |
| 145Nd | GR1 | 154Sm | CD62L | 163Dy | CD25 | 172Yb | CD279 |  |  |
| 146Nd | TER119 | 155Gd | CD80 | 164Dy | RORgt(Intra) | 173Yb | CD115 |  |  |

**Table S3.**
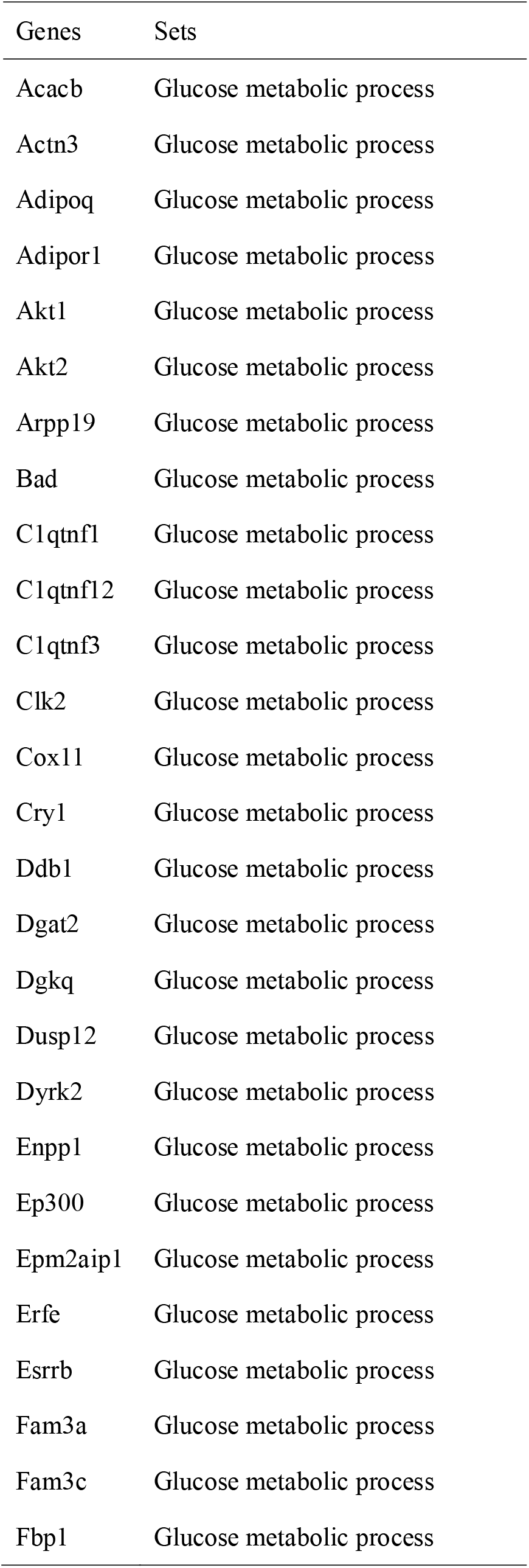
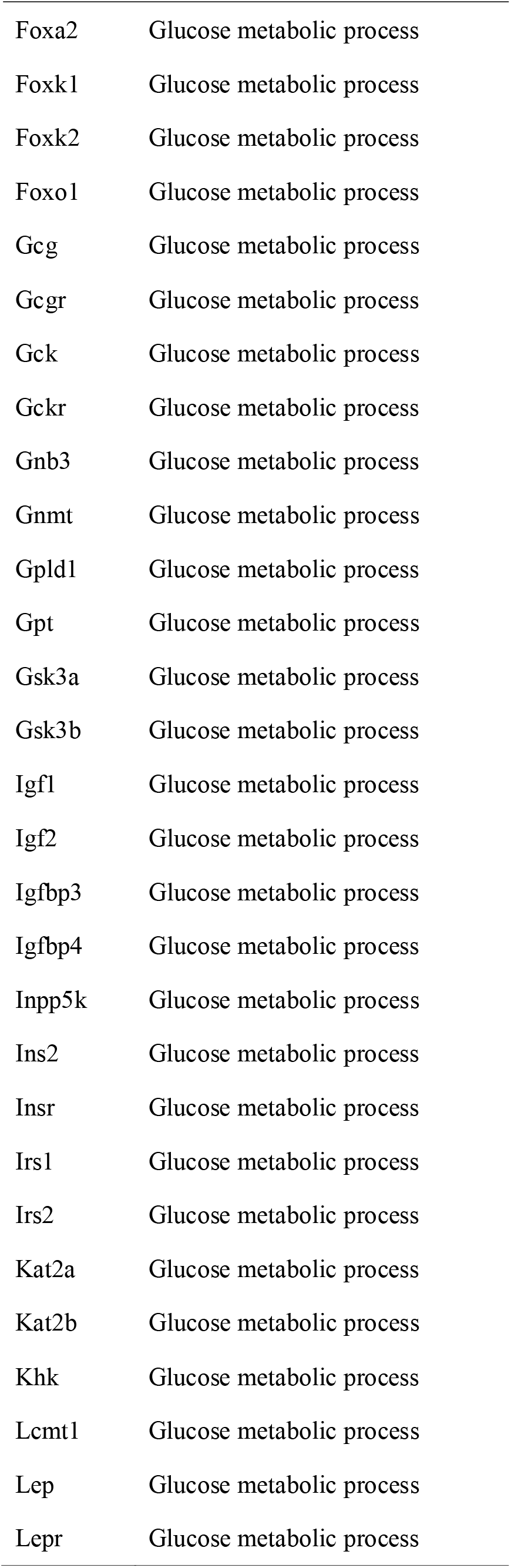
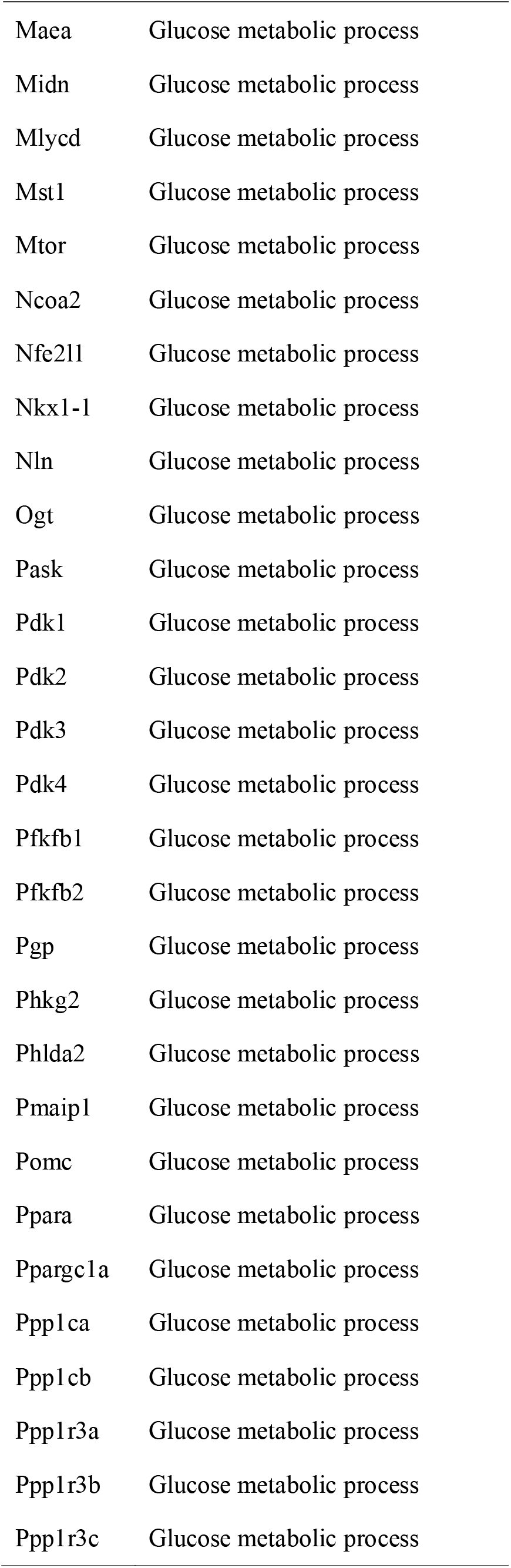
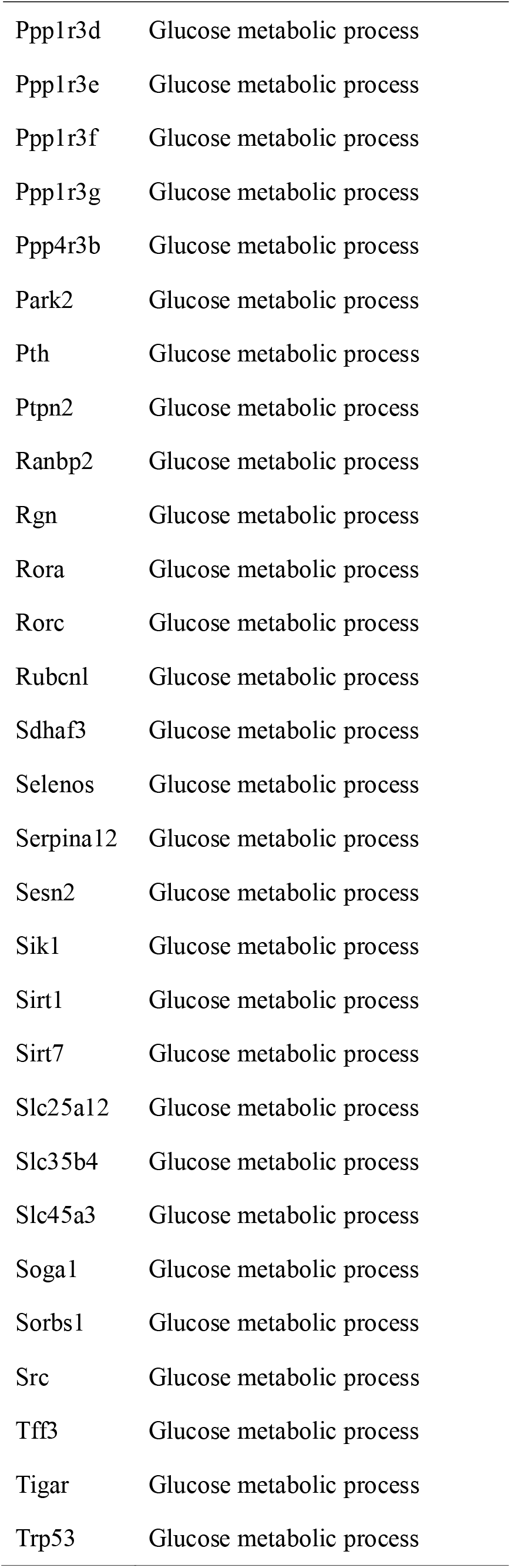
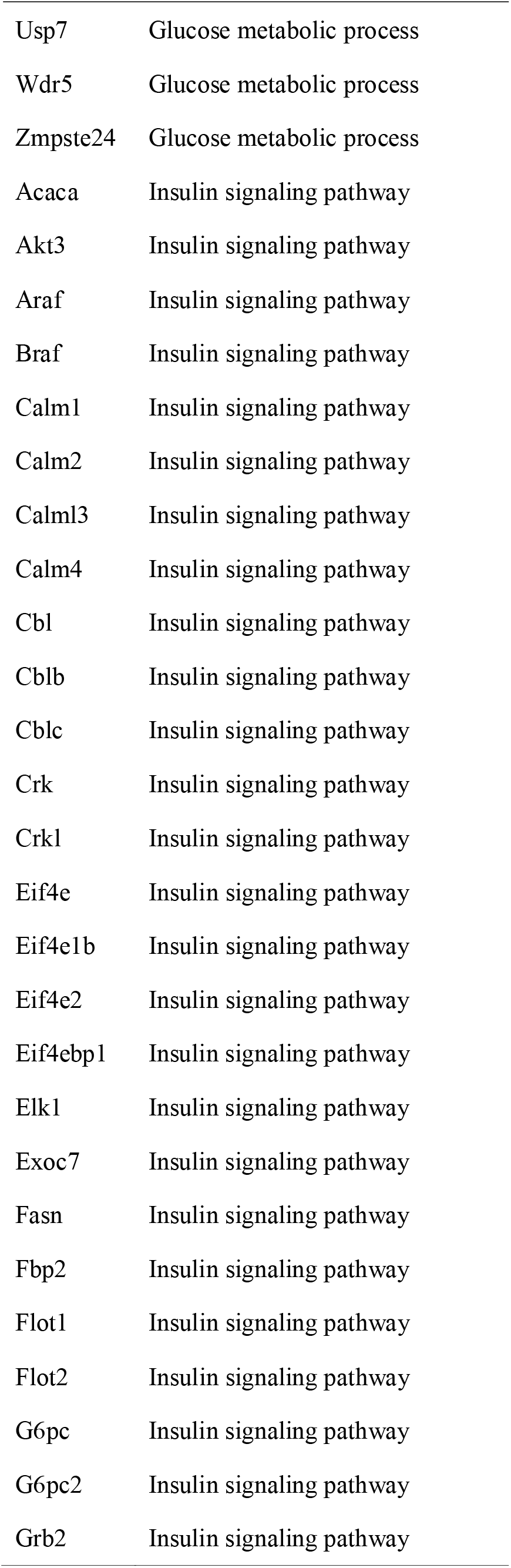
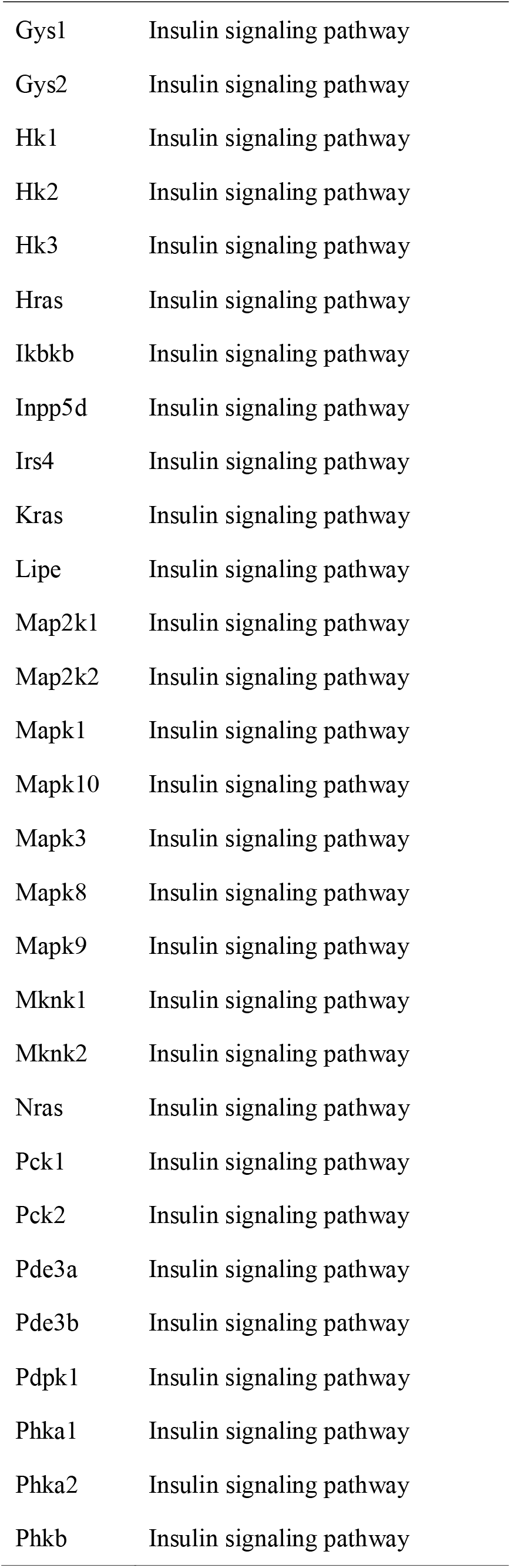
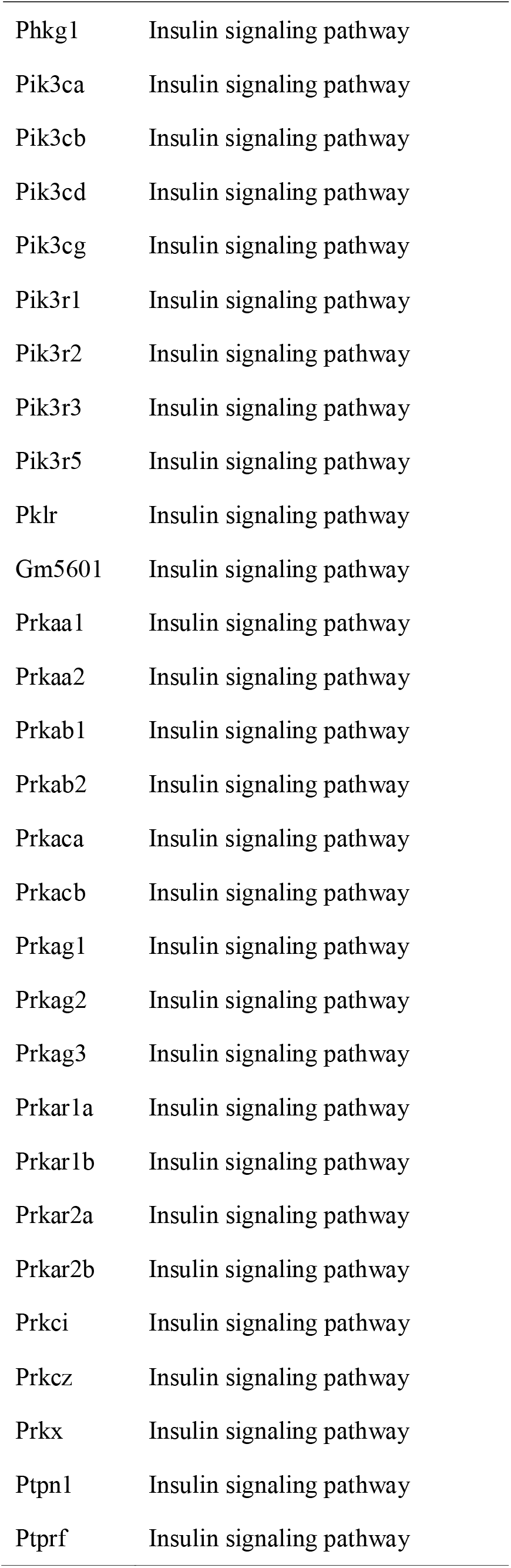
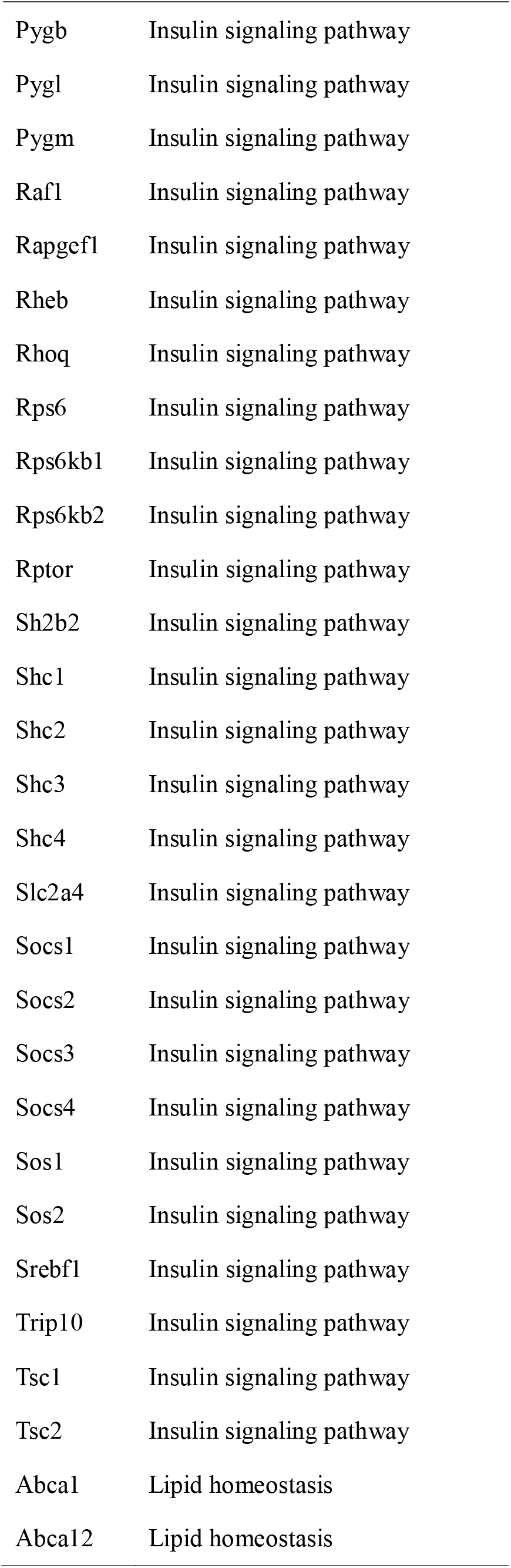
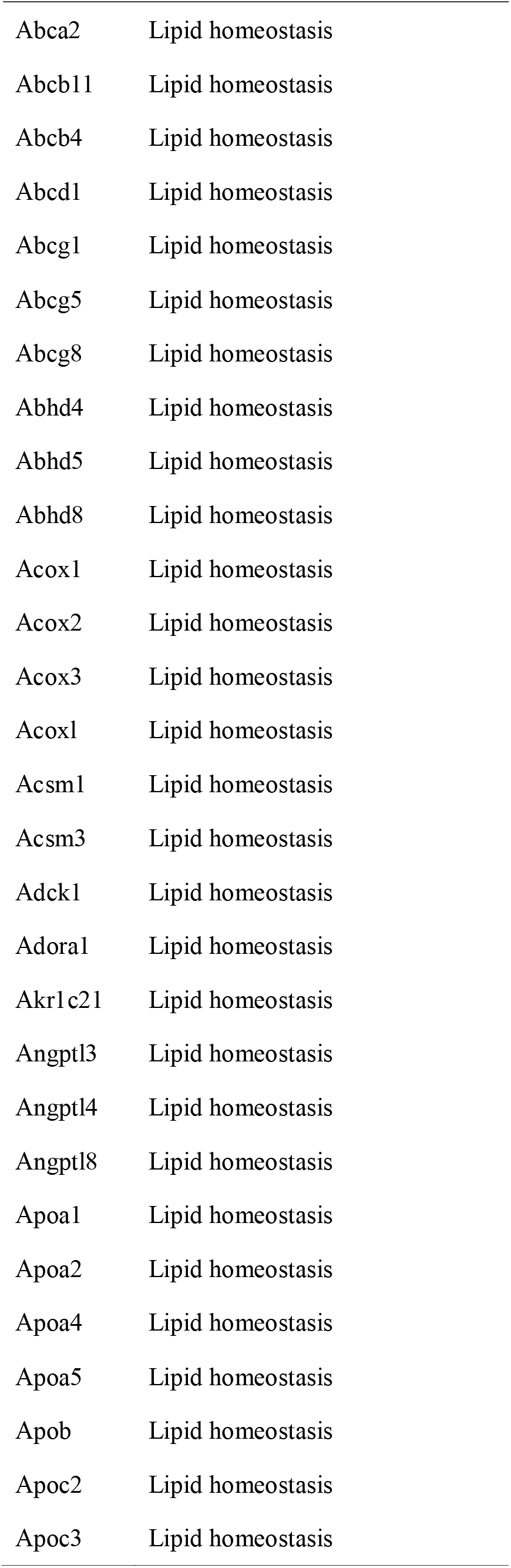
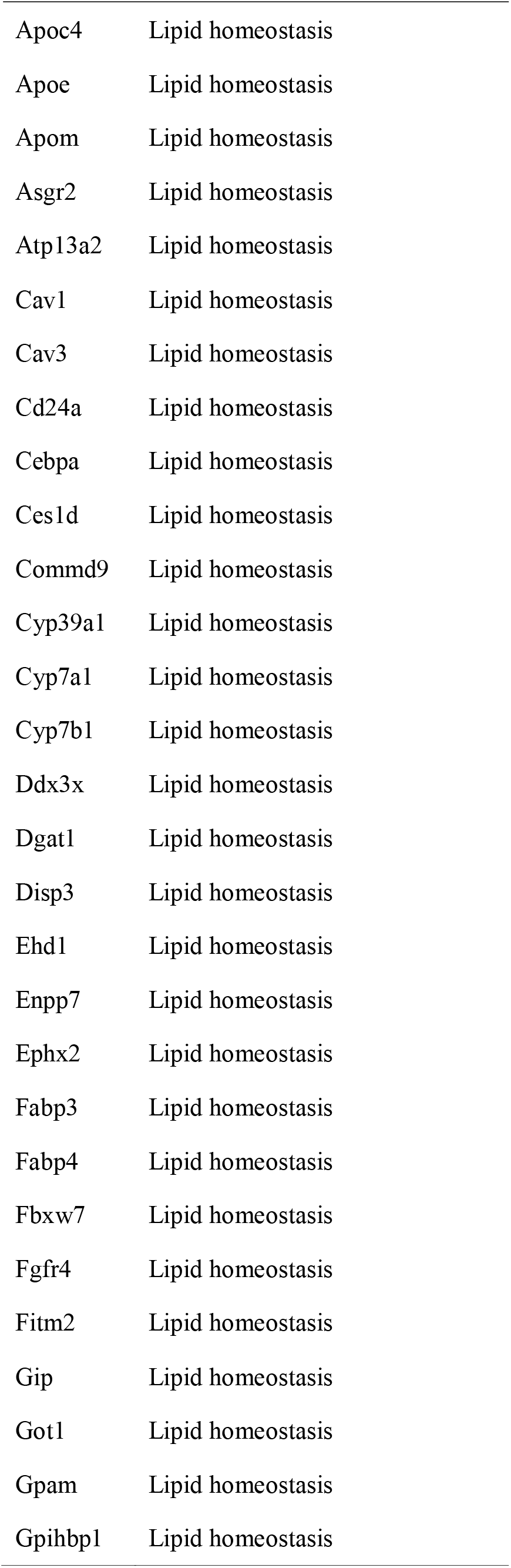
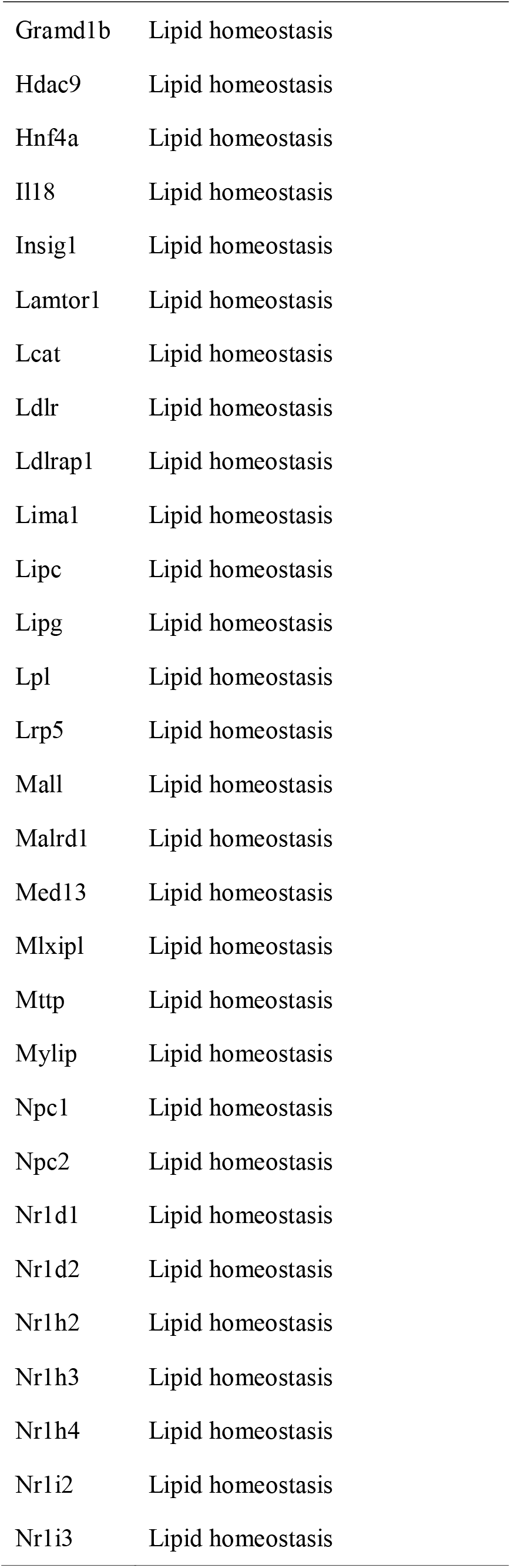
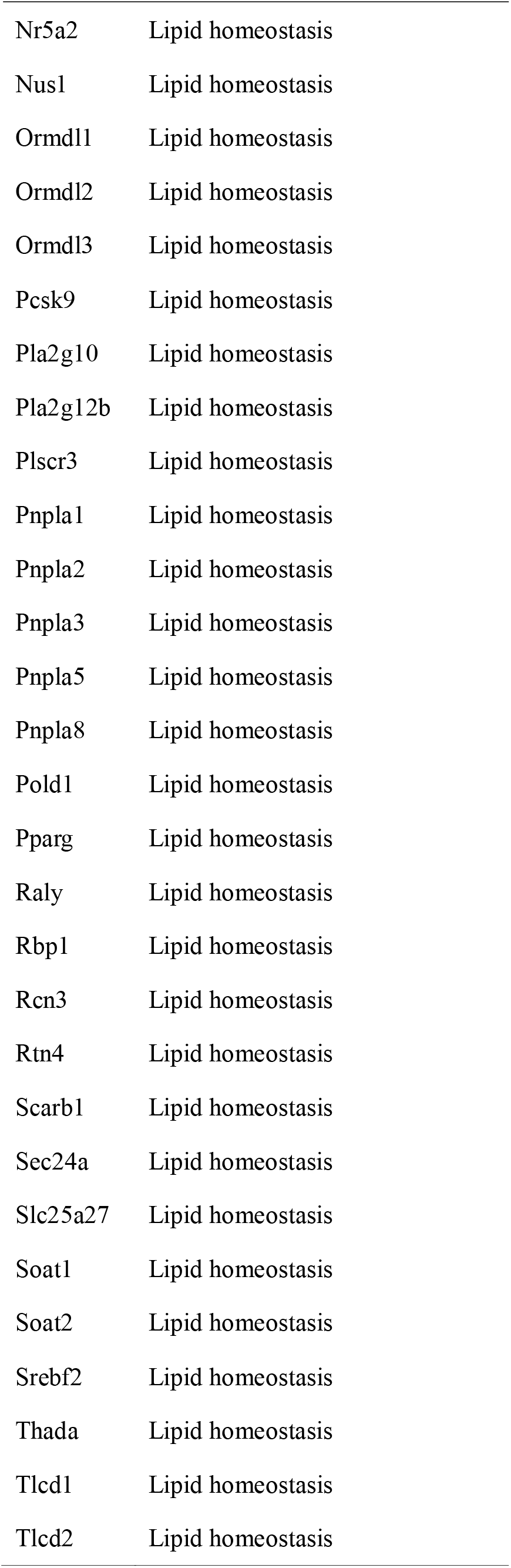
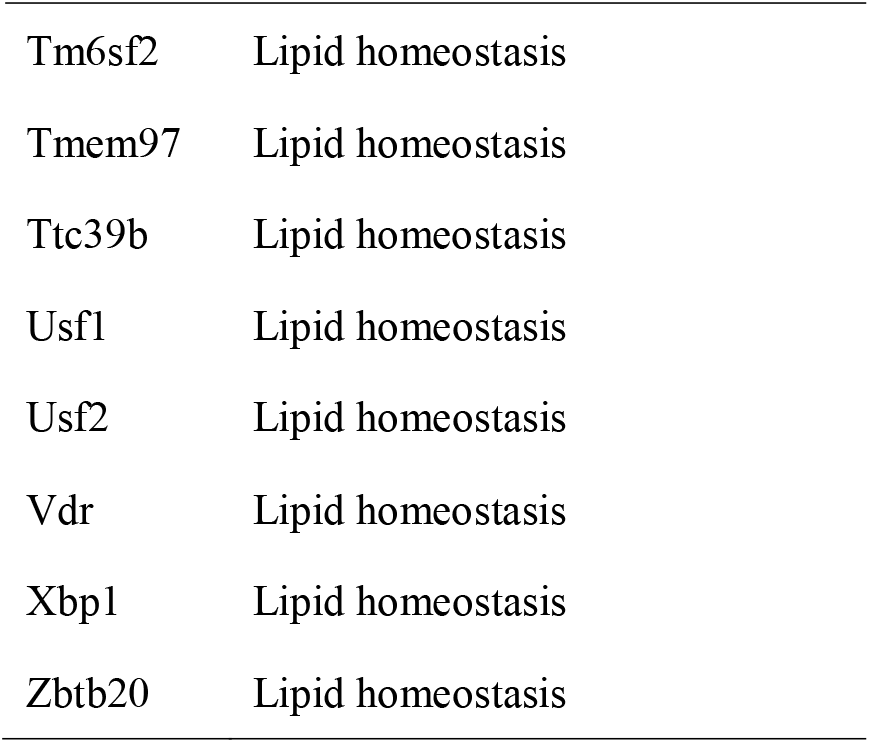
Gene sets of interest.

## REFERENCE

1. Mathur, A.K. and J. Talwalkar, Quality measurement and improvement in liver transplantation. J Hepatol, 2018. 68(6): p. 1300–1310.

2. Lee, D.W. and J. Han, Endoscopic management of anastomotic stricture after living-donor liver transplantation. Korean J Intern Med, 2019. 34(2): p. 261–268.

3. Parmar, A., S.M. Vandriel, and V.L. Ng, Health-related quality of life after pediatric liver transplantation: A systematic review. Liver Transpl, 2017. 23(3): p. 361–374.

4. Yang, L.S., et al., Liver transplantation: a systematic review of long-term quality of life. Liver Int, 2014. 34(9): p. 1298–313.

5. Ling, Q., et al., Association between donor and recipient TCF7L2 gene polymorphisms and the risk of new-onset diabetes mellitus after liver transplantation in a Han Chinese population. J Hepatol, 2013. 58(2): p. 271–7.

6. Bernasconi, E., et al., Airway Microbiota Determines Innate Cell Inflammatory or Tissue Remodeling Profiles in Lung Transplantation. Am J Respir Crit Care Med, 2016. 194(10): p. 1252–1263.

7. Leithead, J.A. and J.W. Ferguson, Chronic kidney disease after liver transplantation. J Hepatol, 2015. 62(1): p. 243–4.

8. Oldani, G., et al., Chimeric liver transplantation reveals interspecific graft remodelling. J Hepatol, 2018. 69(5): p. 1025–1036.

9. Obara, H., et al., IFN-gamma, produced by NK cells that infiltrate liver allografts early after transplantation, links the innate and adaptive immune responses. Am J Transplant, 2005. 5(9): p. 2094–103.

10. Mossanen, J.C., et al., Chemokine (C-C motif) receptor 2-positive monocytes aggravate the early phase of acetaminophen-induced acute liver injury. Hepatology, 2016. 64(5): p. 1667–1682.

11. Song, J.L., et al., A new index predicts early allograft dysfunction following living donor liver transplantation: A propensity score analysis. Dig Liver Dis, 2017. 49(11): p. 1225–1232.

12. Kim, N., et al., Combined Detection of Serum IL-10, IL-17, and CXCL10 Predicts Acute Rejection Following Adult Liver Transplantation. Mol Cells, 2016. 39(8): p. 639–44.

13. Sharma, A., et al., Onco-fetal Reprogramming of Endothelial Cells Drives Immunosuppressive Macrophages in Hepatocellular Carcinoma. Cell, 2020. 183(2): p. 377–394 e21.

14. Lotto, J., et al., Single-Cell Transcriptomics Reveals Early Emergence of Liver Parenchymal and Non-parenchymal Cell Lineages. Cell, 2020. 183(3): p. 702–716 e14.

15. Vento-Tormo, R., et al., Single-cell reconstruction of the early maternal-fetal interface in humans. Nature, 2018. 563(7731): p. 347–353.

16. Browaeys, R., W. Saelens, and Y. Saeys, NicheNet: modeling intercellular communication by linking ligands to target genes. Nat Methods, 2020. 17(2): p. 159–162.

17. Huang, H., et al., The time-dependent shift in the hepatic graft and recipient macrophage pool following liver transplantation. Cell Mol Immunol, 2020. 17(4): p. 412–414.

18. Zhang, C., et al., A Soluble Form of P Selectin Glycoprotein Ligand 1 Requires Signaling by Nuclear Factor Erythroid 2-Related Factor 2 to Protect Liver Transplant Endothelial Cells Against Ischemia-Reperfusion Injury. Am J Transplant, 2017. 17(6): p. 1462–1475.

19. Ling, Q., et al., The tacrolimus-induced glucose homeostasis imbalance in terms of the liver: From bench to bedside. Am J Transplant, 2020. 20(3): p. 701–713.

20. Zhang, Q., et al., Integrated multiomic analysis reveals comprehensive tumour heterogeneity and novel immunophenotypic classification in hepatocellular carcinomas. Gut, 2019. 68(11): p. 2019–2031.

21. Zhang, M., et al., Single-cell transcriptomic architecture and intercellular crosstalk of human intrahepatic cholangiocarcinoma. J Hepatol, 2020. 73(5): p. 1118–1130.

22. Bonnardel, J., et al., Stellate Cells, Hepatocytes, and Endothelial Cells Imprint the Kupffer Cell Identity on Monocytes Colonizing the Liver Macrophage Niche. Immunity, 2019. 51(4): p. 638–654 e9.

23. Bárcena, C., et al., CD5L is a pleiotropic player in liver fibrosis controlling damage, fibrosis and immune cell content. EBioMedicine, 2019. 43: p. 513–524.

24. Wu, H., et al., TIM-4 interference in Kupffer cells against CCL4-induced liver fibrosis by mediating Akt1/Mitophagy signalling pathway. Cell Prolif, 2020. 53(1): p. e12731.

25. Wu, X., et al., Soluble CLEC2 Extracellular Domain Improves Glucose and Lipid Homeostasis by Regulating Liver Kupffer Cell Polarization. EBioMedicine, 2015. 2(3): p. 214–24.

26. Yang, X., et al., Overexpressed PLTP in macrophage may promote cholesterol accumulation by prolonged endoplasmic reticulum stress. Med Hypotheses, 2017. 98: p. 45–48.

27. Song, Q., et al., Dissecting intratumoral myeloid cell plasticity by single cell RNA-seq. Cancer Med, 2019. 8(6): p. 3072–3085.

28. Ma, K.L., et al., Inflammatory stress exacerbates lipid accumulation in hepatic cells and fatty livers of apolipoprotein E knockout mice. Hepatology, 2008. 48(3): p. 770–81.

29. Dou, L., et al., Mir-338-3p Mediates Tnf-A-Induced Hepatic Insulin Resistance by Targeting PP4r1 to Regulate PP4 Expression. Cell Physiol Biochem, 2017. 41(6): p. 2419–2431.

30. Gilbo, N., et al., Liver graft preconditioning, preservation and reconditioning. Dig Liver Dis, 2016. 48(11): p. 1265–1274.

31. Liu, Y., et al., Activation of YAP attenuates hepatic damage and fibrosis in liver ischemia-reperfusion injury. J Hepatol, 2019. 71(4): p. 719–730.

32. Yazdani, H.O., et al., Exercise training decreases hepatic injury via changes in immune response to liver ischemia/reperfusion in mice. Hepatology, 2020.

33. Patrono, D., et al., Perfusate Analysis During Dual Hypothermic Oxygenated Machine Perfusion of Liver Grafts: Correlations With Donor Factors and Early Outcomes. Transplantation, 2020. 104(9): p. 1929–1942.

34. Xiong, X., et al., Landscape of Intercellular Crosstalk in Healthy and NASH Liver Revealed by Single-Cell Secretome Gene Analysis. Mol Cell, 2019. 75(3): p. 644–660.e5.

35. Robinson, M.W., C. Harmon, and C. O’Farrelly, Liver immunology and its role in inflammation and homeostasis. Cell Mol Immunol, 2016. 13(3): p. 267–76.

36. Chong, A.S., B cells as antigen-presenting cells in transplantation rejection and tolerance. Cell Immunol, 2020. 349: p. 104061.

37. Krenkel, O. and F. Tacke, Liver macrophages in tissue homeostasis and disease. Nat Rev Immunol, 2017. 17(5): p. 306–321.

38. Triantafyllou, E., et al., MerTK expressing hepatic macrophages promote the resolution of inflammation in acute liver failure. Gut, 2018. 67(2): p. 333–347.

39. DeBerge, M., et al., MerTK Cleavage on Resident Cardiac Macrophages Compromises Repair After Myocardial Ischemia Reperfusion Injury. Circ Res, 2017. 121(8): p. 930–940.

40. Kazankov, K., et al., The role of macrophages in nonalcoholic fatty liver disease and nonalcoholic steatohepatitis. Nat Rev Gastroenterol Hepatol, 2018.

41. Sun, H., et al., Accumulation of Tumor-Infiltrating CD49a(+) NK Cells Correlates with Poor Prognosis for Human Hepatocellular Carcinoma. Cancer Immunol Res, 2019. 7(9): p. 1535–1546.

42. Weng, S.Y., et al., IL-4 Receptor Alpha Signaling through Macrophages Differentially Regulates Liver Fibrosis Progression and Reversal. EBioMedicine, 2018. 29: p. 92–103.

43. Harmon, C., et al., Natural Killer Cells and Liver Transplantation: Orchestrators of Rejection or Tolerance? Am J Transplant, 2016. 16(3): p. 751–7.

44. Yeung, O.W., et al., Alternatively activated (M2) macrophages promote tumour growth and invasiveness in hepatocellular carcinoma. J Hepatol, 2015. 62(3): p. 607–16.

45. Ling, Q., et al., New-onset diabetes after liver transplantation: a national report from China Liver Transplant Registry. Liver Int, 2016. 36(5): p. 705–12.

46. Ke, Q.H., et al., New-onset hyperglycemia immediately after liver transplantation: A national survey from China Liver Transplant Registry. Hepatobiliary Pancreat Dis Int, 2018. 17(4): p. 310–315.

47. MacParland, S.A., et al., Single cell RNA sequencing of human liver reveals distinct intrahepatic macrophage populations. Nat Commun, 2018. 9(1): p. 4383.

48. Han, X., et al., Mapping the Mouse Cell Atlas by Microwell-Seq. Cell, 2018. 172(5): p. 1091–1107.e17.

49. Tosello-Trampont, A.C., et al., Kuppfer cells trigger nonalcoholic steatohepatitis development in diet-induced mouse model through tumor necrosis factor-α production. J Biol Chem, 2012. 287(48): p. 40161–72.

50. Marais, A.D., Apolipoprotein E in lipoprotein metabolism, health and cardiovascular disease. Pathology, 2019. 51(2): p. 165–176.

51. Koyama, Y. and D.A. Brenner, Liver inflammation and fibrosis. J Clin Invest, 2017. 127(1): p. 55–64.

