## Supplementary figures and images for "Decoding the single-cell landscape and intercellular crosstalk in the transplanted liver: a 4-dimension mouse model"

### Fig.S1

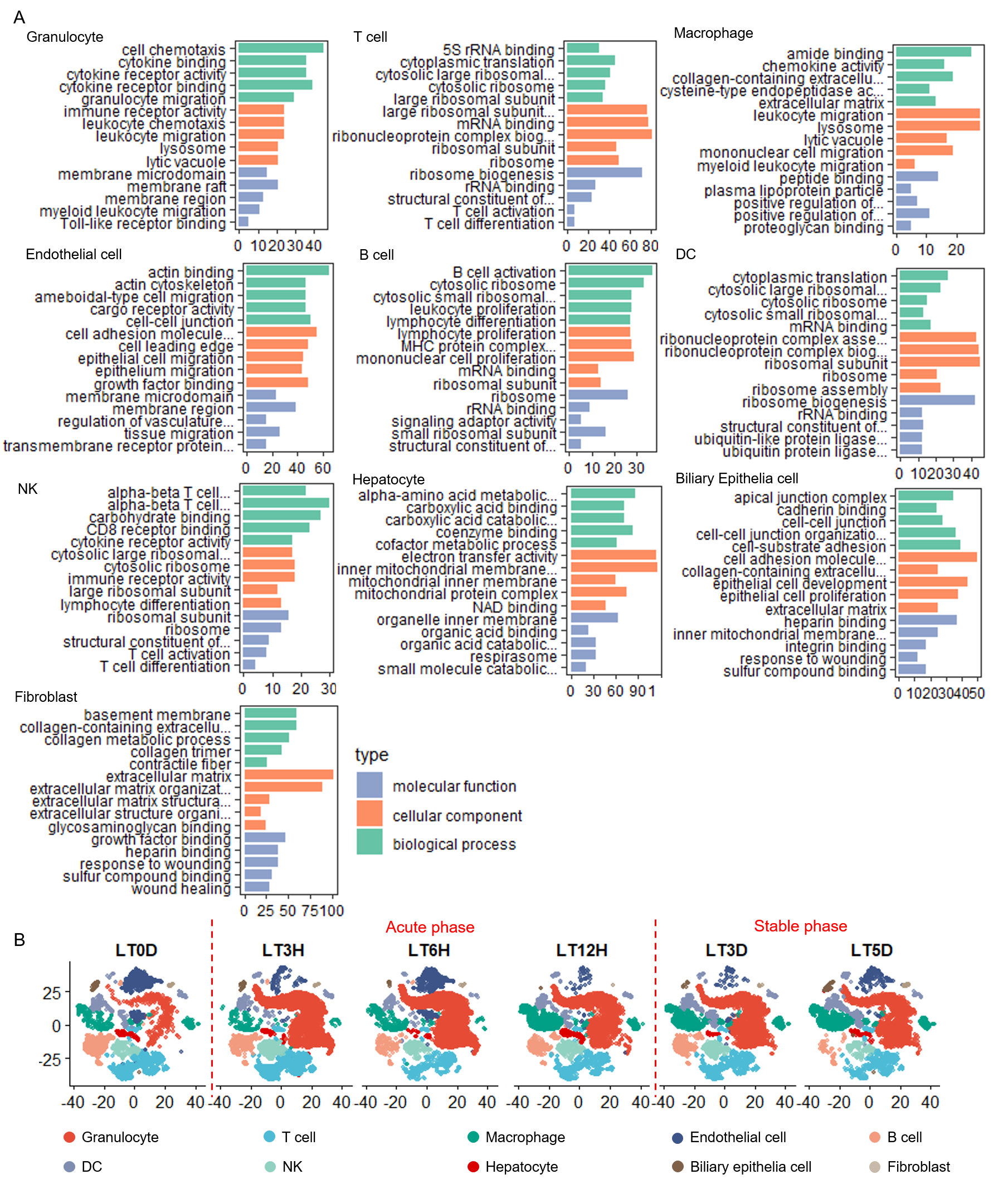
